# Excitatory neurons derived from human induced Pluripotent Stem Cells show transcriptomic differences in Alzheimer’s patients from controls

**DOI:** 10.1101/2023.06.10.544465

**Authors:** Ram Sagar, Ioannis Azoidis, Cristina Zivko, Ariadni Xydia, Esther Oh, Paul Rosenberg, Constantine G. Lyketsos, Vasiliki Mahairaki, Dimitrios Avramopoulos

## Abstract

The recent advances in creating pluripotent stem cells from somatic cells and differentiating them into a variety of cell types is allowing us to study them without the caveats associated with disease related changes. We have generated induced Pluripotent Stem Cells (iPSCs) from eight Alzheimer’s disease (AD) patients and six controls and used lentiviral delivery to differentiate them into excitatory glutamatergic neurons. We have performed RNA sequencing on these neurons and compared the Alzheimer’s and control transcriptomes. We find that 621 genes show differences in expression levels at adjusted p<0.05 between the case and control derived neurons. These genes show significant overlap and direction concordance with genes reported from a Single cell transcriptome study of Alzheimer’s patients, they contain 5 genes implicated with AD from genome wide association studies and they appear to be part of a larger functional network as indicated by an excess of interactions between them observed in the protein-protein interaction database STRING. Exploratory analysis with Uniform Manifold Approximation and Projection (UMAP) suggests distinct clusters of patients, based on gene expression, who maybe clinically different. If confirmed this finding will to contribute to precision medicine approaches to subgroup Alzheimer’s disease.

## INTRODUCTION

Alzheimer’s disease (AD) and dementia globally affect more than 55 million people, and the cases are projected to triple by 2050^1^, causing a major and worsening public health crisis.

Lacking a clinically effective treatment for AD it is important to continue to improve our understanding of its biology including both genetic and environmental contributions. Despite significant progress in the field of AD genetics, including the identification of three genes (PSEN1, PSEN2, and APP) ^2-4^ that cause familial AD, the discovery of APOE as a major risk factor^5, 6^, and the identification of over 30 other genes with smaller but significant contributions^7 8, 9^, the etiological treatments currently approved have not shown substantial improvements in managing the progress of the disease. A promising solution to this problem may be the application of precision medicine to allow a better match between patients and candidate treatments. At the Johns Hopkins Richman Family Precision Medicine Center of Excellence in Alzheimer’s Disease (JH-AD-PMC), we are continuously looking for new ways to achieve this.

One of the most promising ways to accurately characterize patients in order to identify biomarkers that can help towards the goal of precision medicine is the study of their biological materials. Among the samples and materials most likely to provide answers are the cells directly involved in the disease under study. For diseases of the human brain such as AD this poses a significant problem, as there are multiple limitations in the study of brain cells. The brain is not accessible and cannot be sampled without risk. Even if sampled, a mix of cells would be acquired (neurons, astrocytes, microglia etc.) at varying relative abundance. Further, these cells would have been subjected not only to the ongoing disease process, but also to the medications used against it and the many unknown environmental variables. Recent advances in cell engineering have opened new possibilities for such studies. Accessible cells such as skin fibroblasts or peripheral blood mononuclear cells (PMNCs) can be easily acquired and then reprogramed into iPSCs, erasing the epigenetic marks of differentiation and allowing new differentiation to a variety of cell types ^10-13^. Induced PSCs were first established more than a decade ago using the four Yamanaka transcription factors (OCT3/4, SOX2, KLF4, c-MYC) to reprogram mouse adult fibroblast cells^14^. Since then, different technological advances have contributed to reliably establish human iPSCs from different cells, including new combinations of transcription factors, using small molecules, or reprogramming with episomal vectors^15^. Advances in differentiation methods have allowed the generation of multiple cell types from iPSCs, including many types of neuronal and glial cells ^16, 17^ that resemble their in vivo counterparts and have become a popular tool for preclinical disease modeling in scientific research ^18 16^. At the same time, advances in DNA sequencing allow the accurate characterization of the transcriptomic state of cells and tissues, a window into their metabolic state and genomic make-up.

Here we investigate whether cells derived from AD patients show transcriptomic differences as compared to cells derived from healthy individuals, after they have been first reprogrammed to pluripotency followed by differentiation into excitatory neurons. Such differences could not only be useful in predicting disease, but also in categorizing patients into different evidence-based clusters that might predict their course and response to treatment, a big step forward for precision medicine in AD.

## MATERIALS AND METHODS

### Patients and controls

Blood from patients (JHU-AD-01 to -06 and JHU-AD-08), as well as from healthy individuals (JHU-WT-03, -04, -07 and -08) was collected through the Johns Hopkins Alzheimer’s Disease Research Center (ADRC) and the Johns Hopkins Memory and Alzheimer’s Treatment Center (MATC). Blood from patient JHU-AD-07 was provided to us from collaborators through the ongoing S-CitAD clinical trial (ClinicalTrials.gov Identifier: NCT03108846). Patients recruited in this study have AD dementia, Mini-Mental State Examination (MMSE) 5-26 and meet criteria for agitation syndrome ^19^. Adult skin fibroblasts from clinically normal individuals (JHU-WT-01 and -05) were obtained from the Coriell Institute for Medical Research ^20^. AD patients had a mean age of 71.4 years and controls a mean age of 71.7 years. Table 1 shows the age and sex of all subjects. The samples were overwhelmingly female with only 3 males among the patients. Blood was collected by venipuncture into yellow-top tubes (containing trisodium citrate, citric acid and dextrose) and shipped to the Johns Hopkins Genetics Core facility for the isolation of PMNCs where it was stored in liquid nitrogen until used.

**Table 1.**
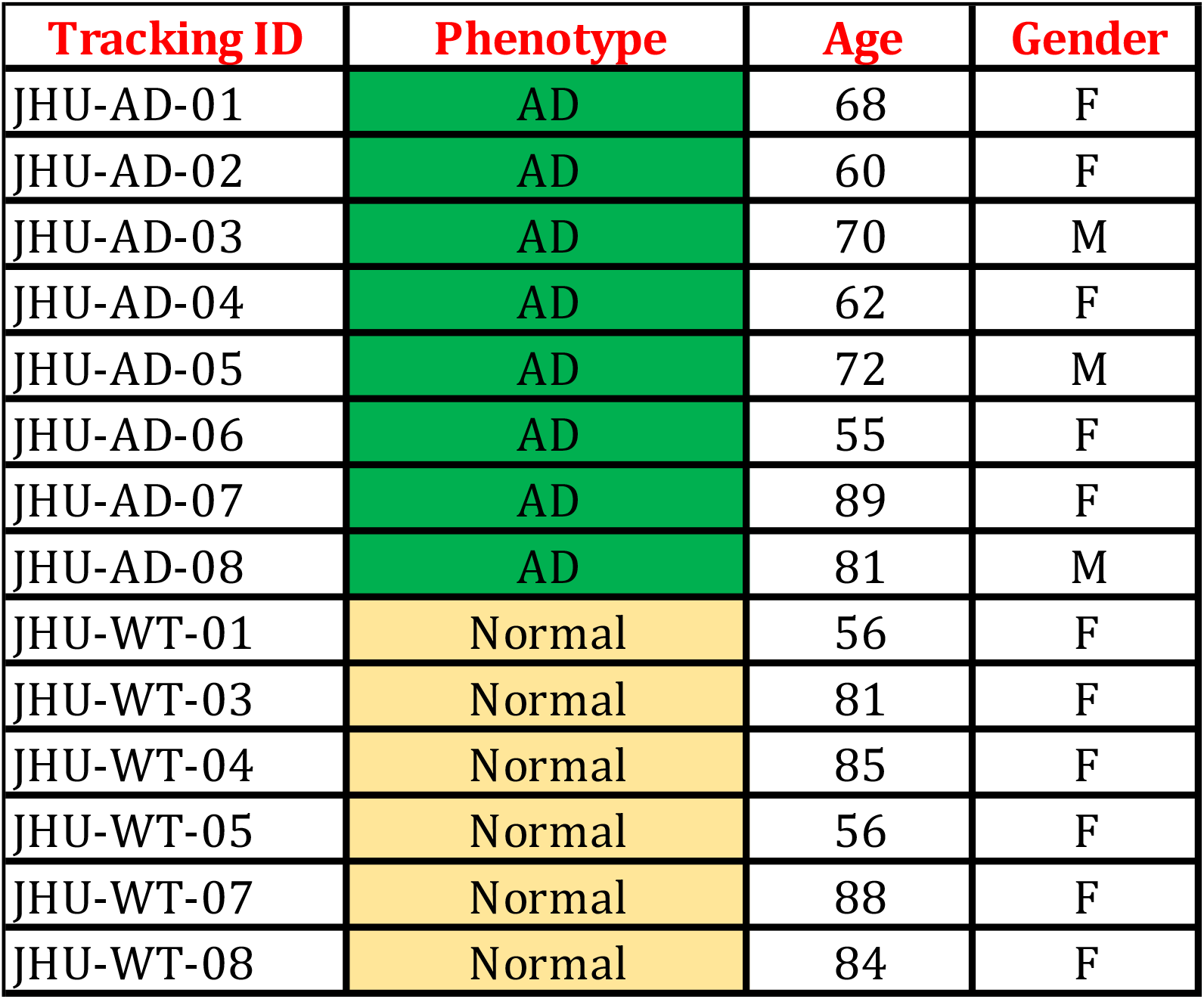
Diagnosis and demographics of patients and healthy individuals used in the transcriptomic studies.

### Induced pluripotent stem cell (iPSC) generation, culture and maintenance

Peripheral blood mononuclear cells (PMNCs) were isolated from the blood samples of the individuals who consented to their participation in the study. They were then reprogrammed to iPSCs by using a transient expression method (nucleofection) involving three plasmid vectors (MOS, MMK and GBX) under feeder-free/xeno-free culture on 4D Nucleofector (Lonza)^11, 21-23^. Generated iPSCs were characterized by immunocytochemistry for pluripotency markers, including Nanog, OCT4 and TRA-1-60, flow cytometry and karyotyping. Established cell lines were cultured (250K cells/well in 6 well plate) on vitronectin-coated tissue culture plates in Essential-8 (E8) medium with 10uM rock inhibitor (Y-27632) during seeding and then maintained in E8 medium till 80-90% confluency ^11, 22^.

### Lentivirus transduction of iPSCs with NgN2

To acquire a high yield of functional neurons we transduced the generated human iPSCs with Ngn2 and rTTA expressing lentivirus (lentivirus was purchased by Cellomics Technology, Maryland USA). We followed an established protocol for generation of induced neuronal cells in 21 days ^17, 24^. Once we established iPSCs confluency at 40-50%, the cells were transduced with Ngn2 (1.5 uL) and rtTA (1.5 uL) expressing lentivirus. Further polybrene (1ug/mL, Santa Cruz) was added at the time of transduction and cells were incubated for 6 hours at 37 ^0^C. After 6 hours, culture medium was replaced with fresh E8 medium and cells were incubated for 24 hours or until 90% confluency was achieved. Cells were hereafter passaged only in rock inhibitor supplemented E8 medium. Transduced cells (250k cells/well in a 6-well plate) were used for neuronal differentiation in this study.

### Neuronal differentiation of Ngn2 transduced iPSCs

The iPSCs were differentiated into induced neuronal cells as described in a previously published protocol ^17, 24, 25^. Ngn2 transduced cells (250K cells/well) were plated on vitronectin coated 6 well culture plates. After 48 hours, culture medium was replaced with fresh E8 medium. To start differentiation, Doxycycline was added to induce Ngn2 expression (Day 0) using the iN-N2 induction medium {DMEM/F12 (Gibco), N-2 supplement (Gibco), D-Glucose (Sigma), 2-β-mercaptoethanol (Gibco), Primocin (Invivogen), BDNF (10 ng/mL, Peprotech), NT3 (10 ng/mL, Peprotech), Laminin (200 ng/mL, Millipore Sigma) and Doxycycline (2 μg/mL, Sigma)}. On Day 1, the induction medium was supplemented with puromycin (2 μg/mL) for 24 hours for selection of transduced cells. On Day 2, surviving cells were passaged on Matrigel-coated 6 well plates at a concentration of 500K cells/well in neural differentiation medium {Neurobasal medium (Gibco), B27 supplement (Gibco), Glutamax (1% v/v, Gibco), Penicillin-Streptomycin (Pen/Strep, 5000 units/mL and 5000 μg/mL respectively, Gibco), D-Glucose (Sigma), BDNF (10 ng/mL), NT3 (10 ng/mL), Laminin (200 ng/mL) and Doxycycline (2 μg/mL, Sigma)}. On Day 4, 50% of the medium was replaced with Neural maturation medium {Neurobasal medium A (Gibco), B27, Glutamax, Pen/Strep, Glucose Pyruvate mix (1:100, final concentration of 5 mM glucose and 10 mM sodium pyruvate), BDNF (10 ng/mL), NT3 (10 ng/mL), Laminin (200 ng/mL) and Doxycycline (2 μg/mL)} supplemented with 4 μM Cytosineβ-D-arabinofuranoside hydrochloride (Ara-C) to inhibit the non-neuronal cell proliferation. From Day 6, 70% of neuronal maturation medium was changed every other day till Day 12. From Day 15, non-doxycycline supplemented maturation medium was used to replace 50% of the medium every 48 hours thereafter till Day 21. Mature neuronal cells were collected on Day 21 for RNA isolation and transcriptomic analysis.

### Immunocytochemistry staining for pluripotency

Immunocytochemistry staining was performed to confirm the pluripotency by using nucleus markers (OCT4 and NANOG) and for surface marker we have used TRA-1-60 antibodies. We cultured the iPSCs in the 12-well plates (60000-cells/well) for 3 days, then cells were quickly washed PBS and 4% PFA (paraformaldehyde) was used for fixation upto 20 minutes in 20^0^C. Cells were washed with PBS and incubated with blocking buffer with 10% goat serum for 1 hour. Primary antibodies OCT4 (#SC-9081), NANOG (#BD-560482), TRA-1-60 (#MAB4360) were used for staining overnight at 4^0^C. Next day cells were washed with PBS and incubated at 37^0^C with secondary antibodies and counterstained with DAPI for 20 minutes at 4^0^C.

### Flow cytometric

In order to validate the pluripotency of the cells, the flow cytometry analysis was completed on a BD LSR Fortessa Analyzer (BD Biosciences) and data were analyzed by using flowJo software. Induced PSCs were dissociated into single cells with TrypLE (Life technology) and washed with BD-FACS staining buffer (Thermofischer, #00422257). Then cells were resuspended in the BD-FACS staining buffer and labelled through anti-human TRA-1-60 antibody (Millipore) and anti-mouse IgM control, PE conjugated (#IC015P) and incubated at 4^0^C for 40 minutes. After incubation, 2ml of PBS was added in the labelled cells and centrifuged for 5 minutes at 1500 RPM. Finally, a 200uL FACS-buffer was used to resuspend the cell and analyzed in BD-Fortessa analyzer.

### RNA extraction and quality control

Neuronal cell pellets were used to isolate the total RNA using Quick-RNA MiniPrep Kit (Zymo Research #R1054) according to the manufacturer’s protocol. Total RNA was quantified using Nanodrop (Thermo Scientific). Total RNA of final concentration (400ng) was aliquoted and shipped to Novogene (novogene.com) for 150 bp paired-end RNA-sequencing. Once all the samples passed the primary quality control, library preparation was initiated. See the previously published paper from our group for the descriptive methodology of RNA sequencing and analysis ^25, 26^.

### RNA Sequencing and data analysis

Next generation sequencing was outsourced to Novogene Corporation Inc. (Sacramento CA). The twelve libraries passed Novogene’s quality control and were sequenced in one batch. The company returned to us .fastq files including on average 44.9 million reads per sample with a maximum of 52.7 million and a minimum of 39.8 million. In order to analyze the samples, the 150bp pair-end reads were aligned to the human reference genome GRCH38 using the package Hisat2 (version 2.2.1)^27^, then SAMtools (version 1.1.4)^28^ was utilized to produce the corresponding BAM files, and stringtie (version 2.1.7)^29^ was used to assemble RNAseq alignments into potential transcripts and estimate their abundance according to GRCh38 human genome annotations^30^. Subsequently raw counts were computed via the Bioconductor package tximport ^31^. Only transcripts with at least 10 reads across all samples were considered for further analysis. The Bioconductor package DESeq2 was used for deferential gene expression analysis^32^ and adjusted p (Padj) was calculated using the Benjamini-Hochberg adjustment.

### Bioinformatics analysis

To identify possible outliers due to experimental conditions we performed principal component analysis using read counts as input and the ggplot2 and ggfortify R packages. For UMAP analysis we used the R functions prcomp and umap.

To validate the results through comparison with other datasets the Differentially Expressed genes (DEG) were filtered by the adjusted p-values by DEseq2 (FDR 0.05, 0.1 and 0.2 for our dataset) and intersected with the data reported by Mathys et al ^33^ for excitatory cortical neurons (FDR between 5x10^−5^, 5x10^−6^ and 5x10^−7^ for the Mathys dataset due to the larger number of positives).

To confirm that the DEGs were a valid gene set that could potentially indicate functional differences between AD and control-derived neurons, we utilized the STRING database of protein-protein interactions ^34^ (https://string-db.org/) to determine whether these genes exhibited more frequent interactions than predicted by chance. Input genes were those with adjusted p-value < 0.05 of which 485 were recognized by STRING. The default parameters were used in the STRING interface, including a medium interaction confidence (0.4) and restricting to only the query genes without an additional level of interactors, as suggested by the authors for meaningful edge (interaction) enrichment results.

This study has been approved by the Johns Hopkins Institutional Review Board (IRB) under numbers NA_00045104 and IRB00038456.

## RESULTS

### Generation of iPSCs and excitatory neurons

Human iPSCs were generated through PMNCs acquired from the Johns Hopkins Core facility. Cells were reprogrammed through transient expression method by using episomal vector. Nucleofected cells were grown in the culture for a minimum of two weeks to develop iPSC like colonies ^11, 22^. The morphological identity of the iPSCs was captured by brightfield microscopy (Figure 1a). The immunofluorescence staining with nucleic markers (NANOG & OCT4) and surface marker (TRA-1-60) confirmed pluripotency which was also validated by flow cytometry for TRA-1-60 positive cells (Figure 1c). The functional pluripotency of the iPSCs was assessed by the in-vitro trilineage differentiation into three germ layers as previously reported ^21^. We did not observe any chromosomal abnormality by karyotyping.

**Figure 1.**
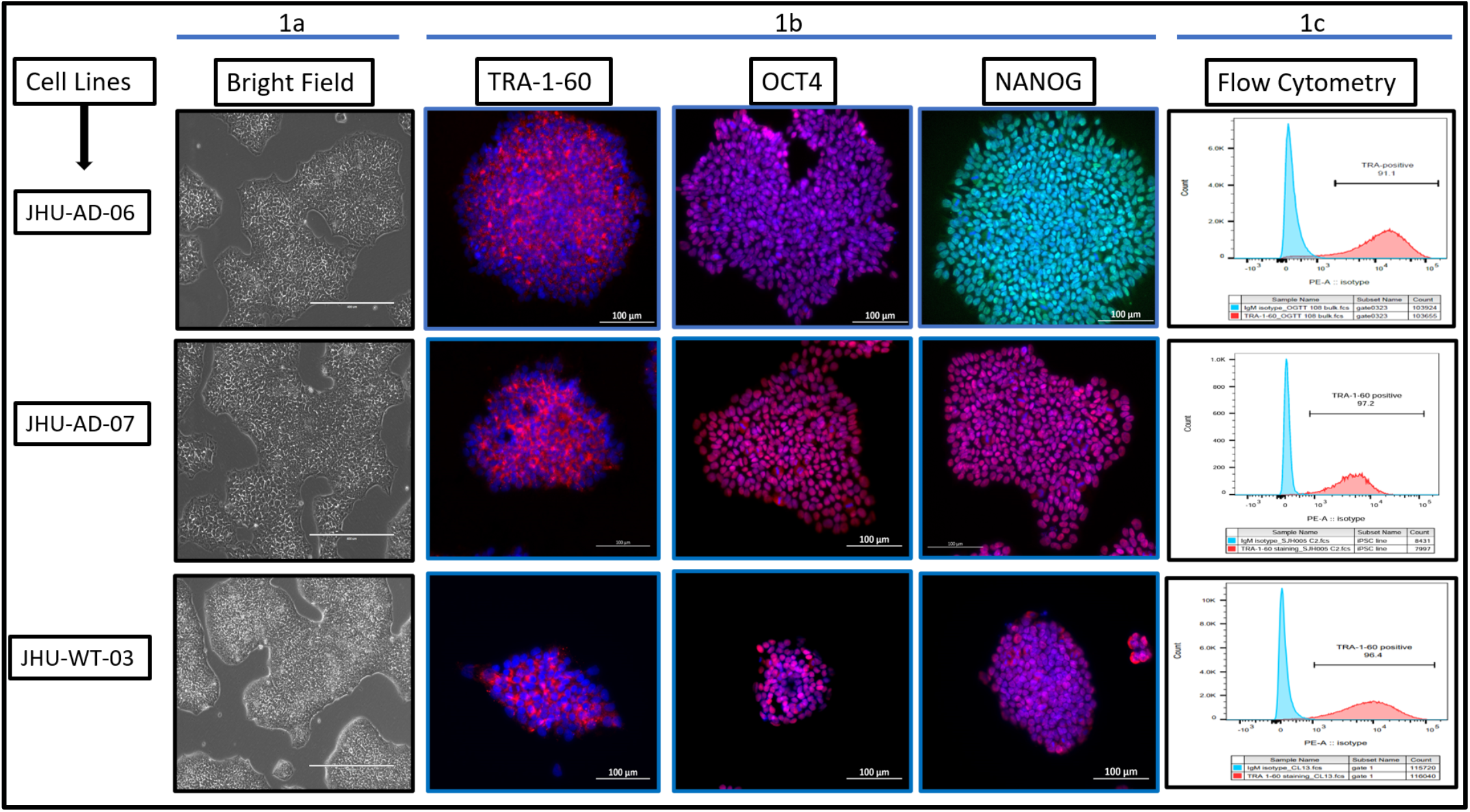
Characterization of iPSCs: **(a)** Brightfield microscope shows the iPSC morphology. **(b)** Fluorescent immunostaining by using pluripotency markers (TRA-1-60, OCT4, Nanog). **(c)** Flow-cytometry shows the purity of the iPSCs by pluripotency marker (TRA-1-60^+^ cells).

To assess the excitatory neuronal identity of the cells produced by Ngn2-induction we used the RNA sequencing results to assess the expression of the expected marker genes. A list of genes and their expression levels is shown in Table 2. We note that we, as well as others, have used this protocol repeatedly and consistently to obtain cells with transcription profiles resembling those of excitatory neurons ^16, 26, 35 24, 25, 36^.

**Table 2.**
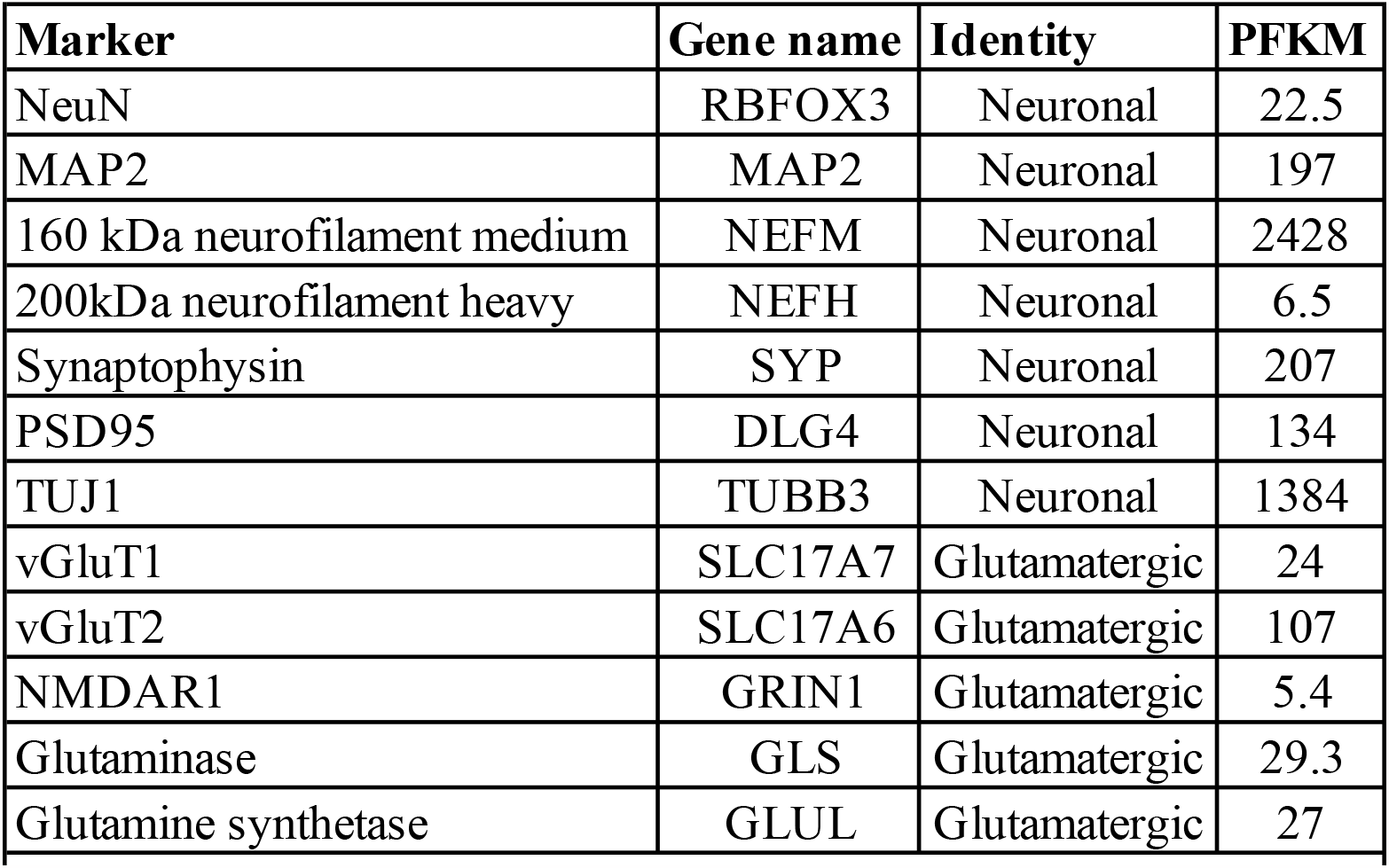
Expression of excitatory neuronal markers in iPSC-derived neurons.

### RNA sequencing

We received on average 45.1 million paired reads per sample (min = 39.8, max = 52.7) with an error rate of 0.03%, of which at least 96.3% had a Phred score of 20 and at least 90.1 of those a Phred score of 30. Principal Component Analysis (PCA) using all genes and the Fragments Per Kbp per million (FPKM) showed that sample AD06 was an outlier, likely the result of unknown differences during cell growth and differentiation, and it was removed from further analysis (Figure 2). The remaining samples were then used for differential expression (DE) analysis. This analysis identified 621/22,011 DE genes at Padj <0.05, 204 higher in AD, and 417 lower.

**Figure 2.**
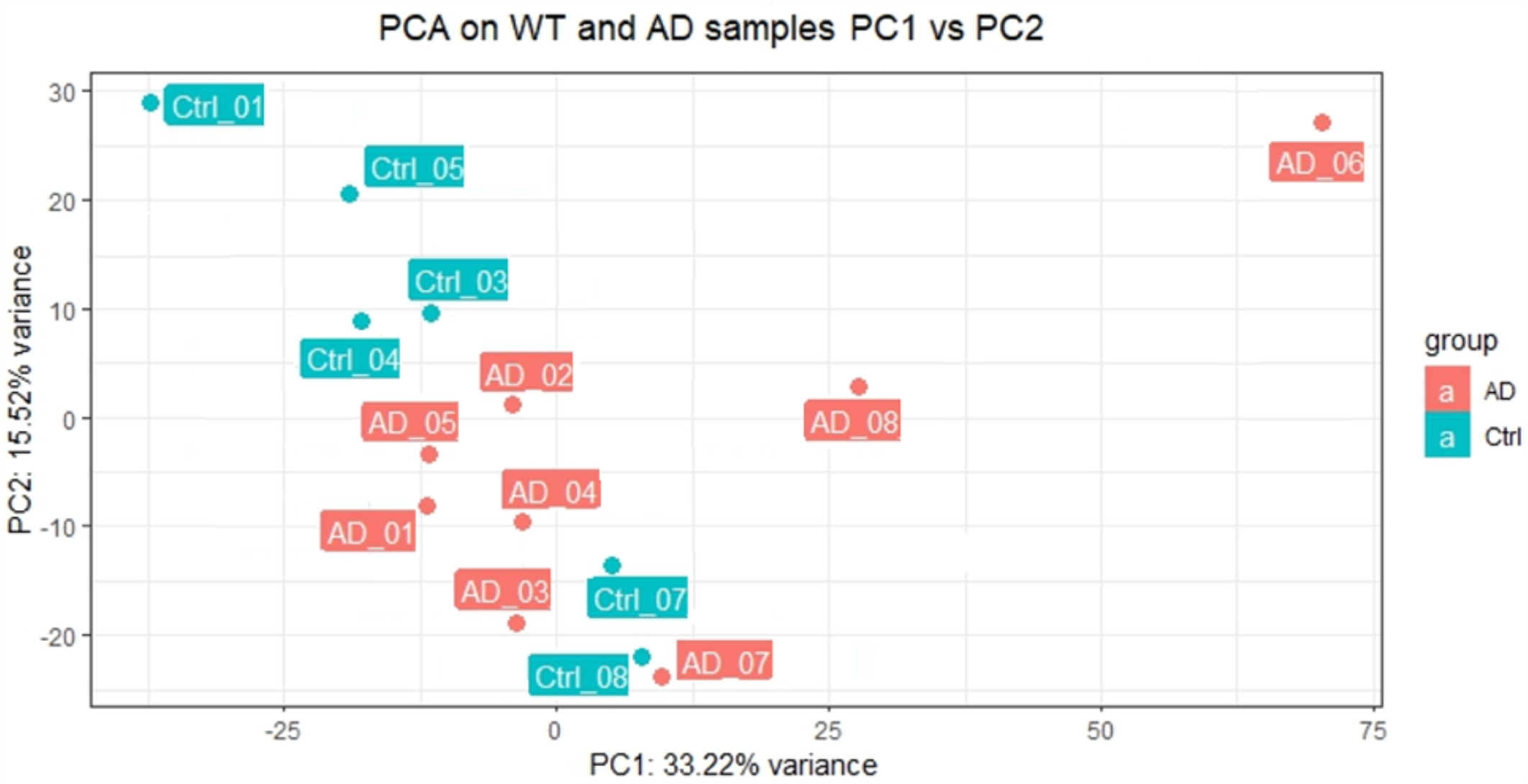
Principal Component Analysis (PCA) using all genes on AD and control samples.

### Bioinformatics analysis

The main question of this project is whether the transcriptome of neuronal cells derived from patients show detectable differences as compare to those derived from controls, making them appropriate for the study of the disease and potentially the identification of biomarkers. Having identified a significant number of genes changing expression, we set out to test whether these genes are a meaningful set and related to AD rather than a collection of random genes from experimental noise. To do so we asked three specific questions: Are the identified genes related to each other? Do they overlap with genes identified by comparable in vivo studies? Do they contain genes that are known to be involved in Alzheimer’s disease? Lastly, we investigated whether the transcriptome of neurons derived from patients and controls could potentially be used for meaningful classification of individuals to facilitate precision medicine.

#### Dysregulated genes are part of a gene network

To explore whether the set of dysregulated genes is an apparently random collection or a functionally meaningful set we queried the STRING database which systematically collects and integrates physical interactions and functional associations. Additionally to the interactive visualization of these interactions in a provided gene set, it allows the assessment of excess of edges (connections) between the nodes (proteins/genes) compared to a random set of the 485 genes corresponding to known proteins at Padj < 0.05 there were 801 interactions, compared to an expected 662 (p-enrichment 9x10-8). The interacting proteins were found to be enriched for those functioning in the brain (FDR=3x10^−5^).

#### Validation through comparison with external data

As opposed to most transcriptomic studies of AD that examine bulk cells from the brain, our data comes from a more homogeneous population of cells after reprogramming and differentiation to excitatory neurons. We sought an appropriate dataset to examine the validity of our results by comparing the identified differentially regulated genes. No published dataset fully matched our conditions, yet one recent study performed single cell sequencing providing data specific to excitatory neurons from the prefrontal cortex of 48 individuals with varying degrees of AD pathology (Mathys et al, ^33^ from their suppl. Table 3, tab Ex, no pathology vs. pathology). Due to our small sample size we examined genes with a corrected p-value of <0.05, 0.1 and 0.2 in our study, while we examined p-value thresholds <5x10^−5^, 5x10^−6^ and 5x10^−7^, for the Mathys et al study which reported many more positive results (Table 2). We consistently found significant excess of overlap from the expected by chance (lowest p= 3.5x10^−5^). We also consistently observed, significantly more concordance in direction of change than that expected by chance (Table 3) reaching a binomial test p-value of 3.2x10^−4^. The enrichment exceeded 1.5-fold and the concordance rate 75% for some thresholds. These results suggest that many of the genes we found to have expression differences mirror changes observed in disease despite our small sample size and despite coming from blood cells that have been reprogrammed and undergone differentiation. It further suggests that the genes driving this overlap represent primary expression changes related to disease risk and not secondary changes due to the disease itself and/or the prescribed medications.

**Table 3.** Comparison of identified differentially regulated genes with external data.

| SC study threshold<br>p (N genes) | Our study threshold p<br>(N genes) | overlap | Fold excess<br>over expected | $\chi^2$ test<br>pvalue | concordance<br>% | concordance<br>N | binomial<br>p-value |
| --- | --- | --- | --- | --- | --- | --- | --- |
| 0.00005 (1331) | 0.05 (394) | 46 | 1.060 | n.s | 72% | 33 | 0.002 |
| 0.000005 (899) | 0.05 (394) | 36 | 1.228 | n.s | 75% | 27 | 0.002 |
| 0.00005 (1331) | 0.1 (765) | 109 | 1.293 | 0.003196507 | 60% | 65 | 0.0275 |
| 0.000005 (899) | 0.1 (765) | 86 | 1.511 | 3.51055E-05 | 62% | 53 | 0.02 |
| 0.00005 (1331) | 0.2 (1547) | 200 | 1.173 | 0.010186373 | 59% | 118 | 0.0065 |
| 0.000005 (899) | 0.2 (1547) | 146 | 1.268 | 0.001361469 | 64% | 94 | 0.00032 |
| 0.0000005 (632) | 0.2 (1547) | 100 | 1.236 | 0.01974401 | 63% | 63 | 0.006 |

**Table 4.**
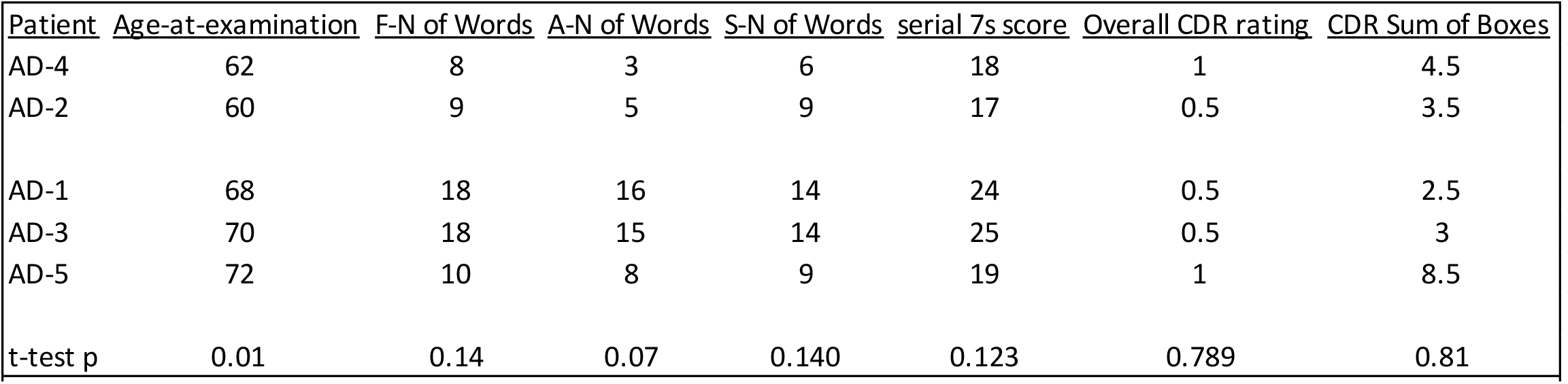
Clinical differences of the identified patient clusters

There were many genes related to AD among the differential expressed (DE) genes between AD and control derived samples. A complete list of genes at all p-values is provided in Supplementary Table 1. Notably, among genes at p-adjusted <0.05 was APOE which was expressed lower in patient-derived neurons and an additional 4 genes identified by large GWAS for AD: *CD2AP, RBFOX1, TMEM132C* and *NPAS2*.

#### Use of transcriptome data as a patient classification tool

Having shown that iPSC-derived neurons from individuals that develop AD show transcriptome differences from controls, we performed exploratory analysis to determine whether the transcriptome of such cells may also be used to cluster affected individuals in groups that might be helpful in determining course and/or best treatment options, a major goal of precision medicine. Given our small sample size we do not expect statistical support, but perhaps an indication on whether this might be a useful path going forward. To determine whether the AD patient transcriptome can be used as a multidimensional variable for patient classification, we used Uniform Manifold Approximation and Projection for Dimension Reduction (UMAP) a technique that can be used to visualize patterns of clustering in high-dimensional data (https://doi.org/10.48550/arXiv.1802.03426) that is frequently used for the analysis of single cell data. Figure 3 shows the results of this analysis. We observed that the 7 patients appeared to form two clusters on the UMAP1 axis, one containing patients 1, 3, 5 and 8 and the other patients 2, 4 and 7. We compared cognitive data between clusters for the individuals with detailed clinical data (all but AD_06 and AD_08). Interestingly, despite having similar clinical dementia ratings (CDR) in the CDR scale ^37^, the two groups were significantly different in age at presentation and the younger group, had consistently lower performance in letter fluency ^38^ “Number of words” test and the mini mental scale ^39^ “Serial 7s” score. These results were not statistically significant, as could be expected by the very small numbers.

**Figure 3.**
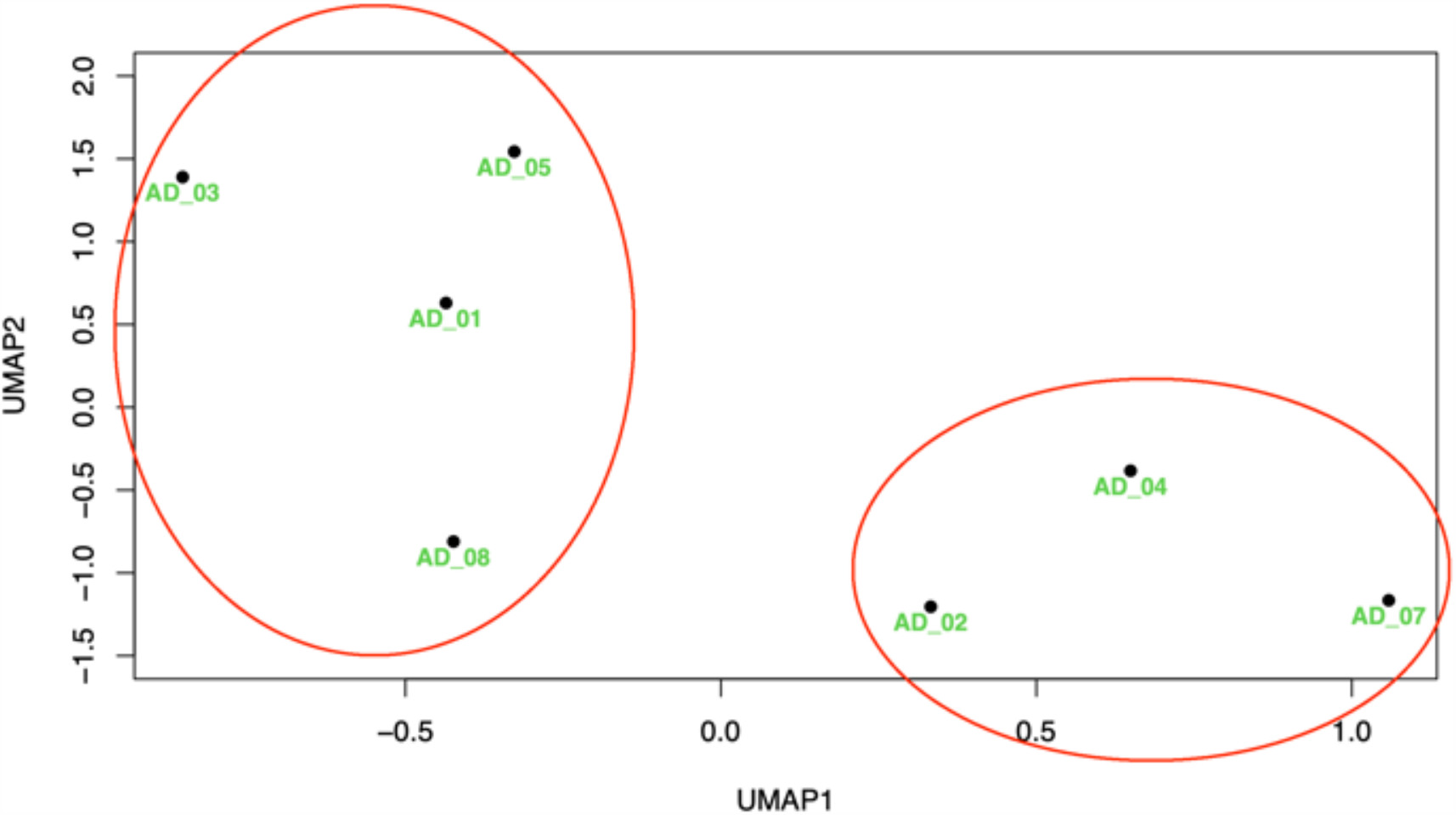
Identification of AD patients’ clusters by UMAP.

## DISCUSSION

We generated iPSCs from 14 individuals and after differentiation to excitatory neurons we found that patients with AD and controls showed significant transcriptomic differences. The set of genes involved appear to be functionally connected to each other and overlap with genes identified by a prior in vivo analysis by another research group, supporting the validity of our results. This is an important observation as it supports the use of measurements from patient-derived cells differentiated into neurons for the identification of biomarkers for AD. Further, our exploratory patient-only analysis for clustering showed two possible clusters, suggesting it might be possible to group patients into biologically meaningful groups that may advance precision medicine.

The genes differing between cases and controls included APOE, a very well established AD risk gene that was found expressed lower in patient derived cells, and others previously linked to AD (*CD2AP, RBFOX1, TMEM132C, NPAS2*). *CD2AP*, whose levels were decreased in the patient derived cells, is a scaffolding molecule reported to be associated with AD ^40^. Its mRNA levels have also been found decreased in peripheral lymphocytes of sporadic AD patients and CD2AP loss of function has been linked to enhanced Aβ production, Tau-induced neurotoxicity, abnormal neurite structure modulation and reduced blood-brain barrier integrity ^41^. *RBFOX1*, whose levels were increased in the patient derived cells is another AD associated gene ^42^, linked to multiple additional psychiatric traits ^43^. It is an RNA-binding protein that regulates alternative splicing ^44^ including that of APP ^45^ and variants near it also regulate gene co-expression modules in the aging human brain ^46^. *TMEM132C*, whose levels were increased in the patient derived cells encodes a neural adhesion molecule associated with AD ^42^ which is also independently associated with cognitive impairment in a hypotensive population ^47^ and with high-altitude adaptation in Tibetans. *NPAS2*, whose levels were decreased in the patient derived cells, is a circadian rhythm gene, a system that has been linked to many diseases including dementia^48^. Interestingly, *NPAS2* has also been implicated in prion diseases ^49, 50^ and is a regulator of genes controlling inflammation in AD ^51^.

While observing biologically relevant differences between the transcriptomes of iPSC-derived neurons from patients and controls, our sample is too small to reach conclusions about possible classification of patients based on these transcriptomes. Our exploratory analysis using UMAP, a tool for visualizing the proximity of different samples in a multidimensional space (transcriptome) in two dimensions suggests two possible clusters. Interestingly the clusters showed some clinical differences, with one cluster (AD_2, and 4) having lower age despite similar clinical dementia ratings and potentially worse scores in specific cognitive tests. We expect that with larger samples more such differences to emerge that might reflect patient subgroups and determine the usefulness of this clustering approach for precision medicine. Incorporated in our mainstream analysis of patients for precision medicine this could become a powerful tool. Our positive results despite the small sample size indicates that the differences we observe are likely only the tip of the iceberg of information we might achieve with large patient samples.

## AKNOWLEGMENTS

This work was supported from The Richman Family Precision Medicine Center of Excellence in Alzheimer’s Disease.

